# OPUS-Rota5: A Highly Accurate Protein Side-chain Modeling Method with 3D-Unet and RotaFormer

**DOI:** 10.1101/2023.10.17.562673

**Authors:** Gang Xu, Zhenwei Luo, Yaming Yan, Qinghua Wang, Jianpeng Ma

## Abstract

Accurate protein side-chain modeling is crucial for protein folding and design. This is particularly true for molecular docking as ligands primarily interact with side chains. A protein structure with large errors in side chains has limited usage such as in drug design. Previous research on AlphaFold2 (AF2) predictions of GPCR targets indicates that the docking of natural ligands back on AF2-predicted structures has limited successful rate presumably due to large errors in side chains. Here, we introduce a two-stage side-chain modeling approach called OPUS-Rota5. It leverages a modified 3D-Unet to capture the local environmental features including ligand information of each residue, and then employs RotaFormer module to aggregate various types of feature. Evaluation on three test sets, including recently released targets from CAMEO and CASP15, reveals that side chains modeled by OPUS-Rota5 are significantly more accurate than those predicted by other methods. We also employ OPUS-Rota5 to refine the side chains of 25 GPCR targets predicted by AF2 and then performed docking of their natural ligands back with a significantly improved successful rate. Such results suggest that OPUS-Rota5 could be a valuable tool for molecular docking, particularly for targets with relatively accurate predicted backbones, but not side chains.

## Introduction

Computational biology is taking on an ever-increasingly important role in the broader field of biology. One critical sub-problem within this domain is protein side-chain modeling, which focuses on reconstructing the side chains for a given protein backbone. This task is vital for protein structure prediction, refinement, and design, as the unique structure of a protein is largely determined by the packing of its side chains^1^. Xu *et al*. ^2^ view protein mutations as side-chain replacements in the corresponding residues and study the effects of these mutations through the lens of side-chain modeling. McPartlon *et al*. ^3^ engage in the co-design of protein sequences using a side-chain modeling framework. In addition, the accurate modeling of side-chain conformations in interfacial residues is crucial for protein-protein interactions ^4, 5^ and molecular docking ^6, 7^. Precise side-chain conformation enhances the accuracy of evaluation of relevant energy functions, which is essential for high-accuracy docking.

Recently, He *et al*. ^6^ examine the molecular docking results on structures predicted by AlphaFold2 ^8^, as well as those on experimental structures of G protein-coupled receptors (GPCRs). Their findings indicate that while AlphaFold2’s predicted backbones closely align with experimental structures, the side chains of certain residues carry significant errors, potentially impacting docking outcomes. Given this, the development of more accurate side-chain modeling methods may be particularly important for small molecule drug screening. This is especially true for target proteins that lack experimentally determined structures but have backbones that are relatively accurately predicted.

Over the past several decades, numerous methods have been proposed for protein side-chain modeling ^1, 3, 9–25^. For instance, in the early 1990s, Holm and colleagues ^24^ developed a database algorithm to generate protein backbone and side-chain coordinates from a C_α_ trace. Meanwhile, Levitt and his team ^25–28^ explored protein folding using molecular dynamics and modeled protein conformation through automatic segment matching. These pioneering studies lay the foundation for the subsequent research in the field.

Sampling-based side-chain modeling methods typically consist of three main components: a rotamer library, a scoring function, and a search algorithm. To minimize the sampling space, some studies have concentrated on developing rotamer libraries that contain a limited set of representative side-chain conformations ^29–32^. A precise scoring function is crucial for effective side-chain packing. In addition to incorporating physics-based energy terms like Van der Waals (vdW) energy and hydrogen bonding energy, some researches have focused on introducing more accurate statistical potentials to improve results ^1, 12, 13, 15^. The efficiency of the search algorithm is vital for the speed of side-chain modeling, most algorithms are based on approaches such as simulated annealing Monte Carlo ^15^, dead-end elimination ^33^ and tree decomposition ^10, 11^.

With the development of deep learning techniques ^34^, several side-chain modeling methods based on deep learning frameworks have emerged. OPUS-Rota3 ^19^ employs a 2D convolutional neural network to extract information from backbone torsion angles, secondary structures, and contact maps, ultimately making dihedral predictions. DLPacker ^17^ and OPUS-Rota4 ^20^ utilize 3D convolutional neural networks to capture the local environmental features of each residue. Additionally, OPUS-Rota4 employs a gradient-based folding framework ^35^ to optimize side-chain conformations using its predicted side-chain contact map. GeoPacker ^14^ and AttnPacker ^3^ take a graph-based approach to capture local features, and they both boast relatively high running speeds. Meanwhile, DiffPack ^21^ employs a torsional diffusion model to autoregressively generate protein side-chain conformations.

In this study, we introduce OPUS-Rota5, a two-stage method for side-chain modeling. The first stage employs a modified 3D-Unet ^36^ to capture the local environmental features of each residue. Inspired by the Evoformer block from AlphaFold2 ^8^, we propose RotaFormer in the second stage to aggregate various types of information, which include 1D protein sequence features, 2D residue contact features, and 3D local environmental features. We assess the performance of OPUS- Rota5 against other leading side-chain modeling methods using three test sets (CAMEO65, CASP15 and CAMEO82). The results indicate that OPUS-Rota5 significantly outperforms these methods. Specifically, on the CASP15 test set, the side-chain RMSD values measured by all residues and core residues are 0.791 and 0.341 for OPUS-Rota5, while they are 0.842 and 0.439 for the second-best method AttnPacker.

Additionally, we use AlphaFold2 to predict the structures of 25 GPCRs and then reconstruct their side chains using OPUS-Rota5. We proceed to perform molecular docking using their natural ligands with AutoDock-Vina ^37, 38^ and Docke6.10 ^39^. Our results demonstrate that the success rates for both docking methods improve after side-chain reconstruction with OPUS-Rota5. For instance, the average RMSD of the top 1 pose generated by AutoDock-Vina and Dock6.10 are 7.577 and 8.548 when tested on the original AlphaFold2 predicted structures. These numbers improve to 6.610 and 7.333 when tested on structures with side chains reconstructed by OPUS- Rota5.

## Method

### Framework of OPUS-Rota5

OPUS-Rota5 employs a two-stage approach for protein side-chain modeling. In the first stage, we leverage a modified 3D-Unet ^36^ to capture each residue’s local environmental features. Drawing inspiration from DLPacker ^17^, we isolate a 20 Å box around each residue to encompass its local environment. We then align the atoms within this box to reference coordinates. Every atom is depicted as a 3D Gaussian density kernel, placed on a 40x40x40 bin grid. We use 28 channels for representation: five for elemental channels (C, N, O, S, and others), one for partial charge (sourced from the Amber99sb force field ^40^), 21 for residue types (20 for common residues and one for others), and one for target residue embedding. The modified 3D-Unet yields two output branches: the first, with four channels (C, N, O, S), provides the 3D side-chain density of the target residue, while the second, with two channels, offers the probability of each bin containing side-chain density. Further architectural details are illustrated in Figure 1B. We then discard the peripheral bins (5 bins from every direction) and down-sample the density map from (40, 40, 40, 4) to (15, 15, 15, 4). Additionally, we incorporate the probability of each bin containing side-chain density as a fifth channel, culminating in the 3D features for each residue. Note that, if we represent the ligand as corresponding atoms in 3D grids, the ligand information can be captured by OPUS-Rota5.

**Figure 1.**
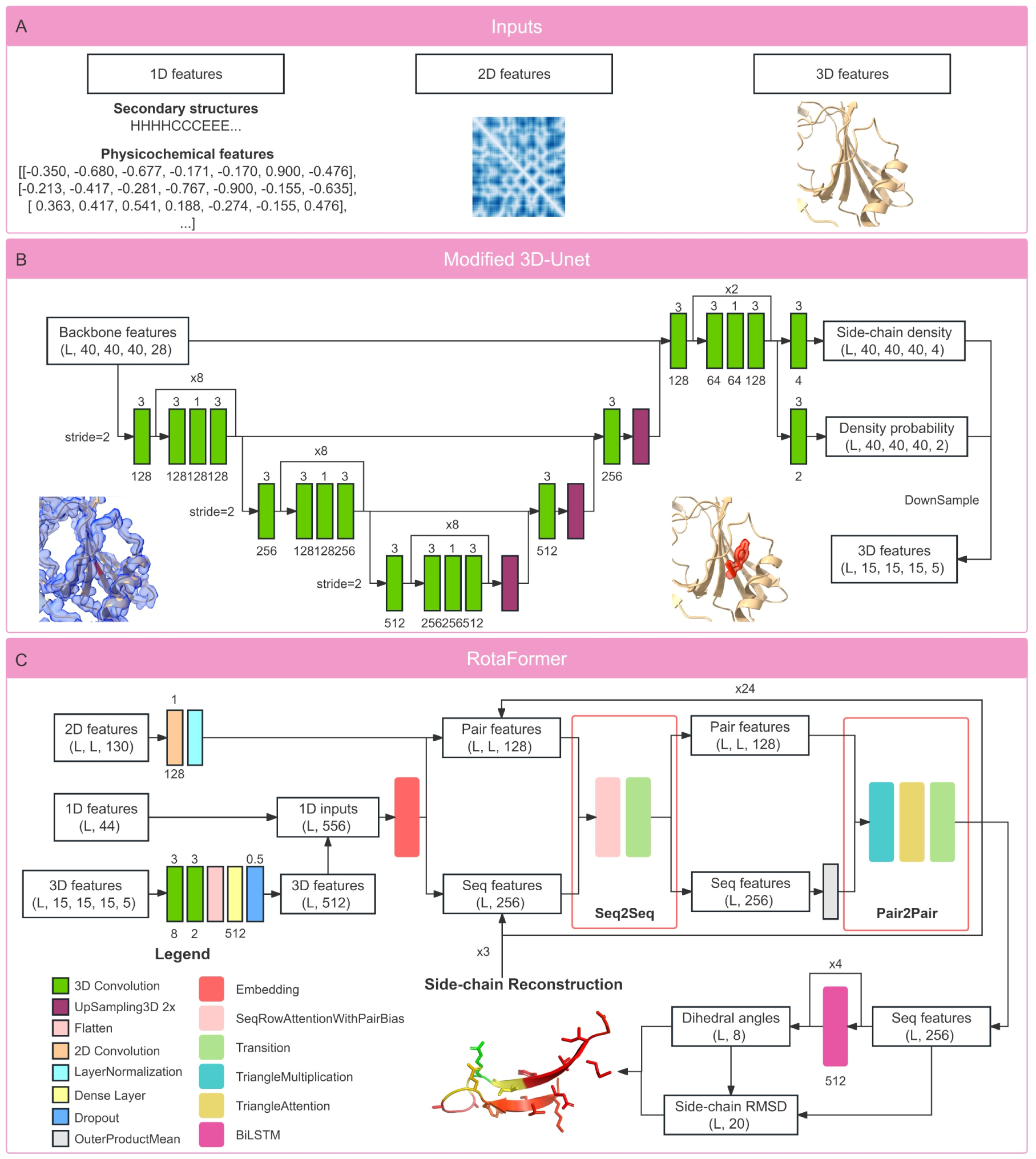
Overview of the OPUS-Rota5 workflow. A. OPUS-Rota5 takes in three primary inputs: 1D protein sequence features, 2D residue-residue contact features, and the 3D backbone density of each residue. B. A modified 3D-Unet is utilized to capture the 3D local environmental features of each residue. C. RotaFormer module is employed to amalgamate various feature types. It produces four predicted side-chain dihedral angles and a predicted side-chain RMSD for each residue. For a given dihedral angle χ, it provides two separate predictions for sin(χ) and cos(χ), respectively.

For training the first branch, we use the mean absolute error (MAE) loss between the predicted and actual side-chain densities. The second branch is trained using cross-entropy loss. We employ the Glorot uniform initializer and the Adam optimizer ^41^. The starting learning rate is set at 1e-3 and is halved if the validation set’s accuracy diminishes. Training ceases after the rate has been reduced four times. We train three models, and the average of their outputs determines the final prediction.

In the second stage, we propose RotaFormer, largely inspired by the Evoformer block in AlphaFold2 ^8^, to consolidate various feature types (as shown in Figure 1A). The 1D protein sequence features encompass the 3-state and 8-state secondary structures, 7 physicochemical properties ^42, 43^, 19 PSP features representing 19 rigid-body blocks within residues ^43–45^, and 6 backbone torsion angle features (comprising sine and cosine values for ϕ, ψ, and ω). Additionally, there’s a confidence feature derived from the DLPacker’s library. The 2D residue-residue backbone contact features, sourced from trRosetta ^46^, include the C_β_- C_β_ distance distributions and the orientational distributions of 3 dihedrals (ω, θ_ab_, θ_ba_) and 2 angles (φ_ab_, φ_ba_) between residues a and b. Distances span from 2 to 20 Å, segmented into 36 bins at 0.5 Å intervals, with an additional bin for distances exceeding 20 Å. The φ angle varies from 0 to 180°, divided into 18 bins at 10° intervals, plus an extra bin for non-contact scenarios. Both ω and θ range from -180 to 180°, segmented into 36 bins at 10° intervals, with another bin for non-contact instances. The 3D local environmental features are the outputs from the modified 3D-Unet in the first stage.

As depicted in Figure 1C, the 3D features undergo two 3D convolution layers. They are then flattened and merged with the 1D features to create the 1D inputs. An embedding module is employed to transform these 1D inputs into sequence and pair features. Concurrently, the 2D features pass through a 2D convolution layer and are added to the pair features. RotaFormer module is adopted to aggregate the sequence and pair features. In particular, it treats the pair features as a bias term, adding it to the attention matrix for each residue pair in the sequence. This effectively integrates the pair features into the sequence features. The module also uses the outer product operation to incorporate sequence features into pair features. We employ 24 RotaFormers for enhanced feature extraction. Subsequently, four BiLSTM layers ^47^ are used to further aggregate sequence features. The output of the second stage is also bifurcated: the first branch provides eight values for side-chain dihedral angle prediction (sine and cosine values for *χ*_1_ to *χ*_4_), while the second branch predicts the root-mean-square deviation (RMSD in Å) of the predicted side chain relative to its native counterpart for each residue. The side-chain RMSD range is set between 0 and 1 and is divided into 20 bins. In the final step, the predicted dihedral angles are integrated back into the initial sequence features, and the aforementioned process is repeated three times.

For training the first branch in RotaFormer module, we introduce the auxiliary loss *L_anglenorm_* from AlphaFold2 ^8^ to ensure the predicted points remain close to the unit circle. For a specific dihedral angle , OPUS-Rota5 predicts values for sin(*χ*) and cos(χ), respectively. The loss function is defined as:

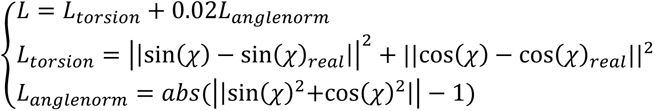

Furthermore, we exclude the loss for dihedral angles not present in certain residues. The second branch, on the other hand, is trained using cross-entropy loss.

Mirroring the training approach of the first stage, we employ the Glorot uniform initializer and the Adam optimizer. Meanwhile, the starting learning rate is set at 1e-4. We also train three models, and the median of their outputs determines the final prediction. Additionally, OPUS-Rota5 is developed in TensorFlow v2.4 ^48^ and trained on a single NVIDIA Tesla V100.

### Datasets

In OPUS-Rota5, we employ the same training dataset as used in OPUS-Rota4 ^20^. This dataset includes 10,024 proteins for training and 983 proteins for validation, all of which were obtained from the PISCES server in February 2017 ^42^.

To assess the side-chain modeling performance across various methods, we utilize three native backbone test sets: CAMEO65 contains 65 hard targets released between May 2021 and October 2021, sourced from the CAMEO website ^49^; CASP15 contains 44 targets, available for download from the CASP website (http://predictioncenter.org); and CAMEO82 contains 82 targets released between May 2023 and August 2023, also from the CAMEO website. Furthermore, we have curated a set of 25 complexes involving G protein-coupled receptors (GPCRs) and their natural small molecule ligands as docking targets.

### Performance Metrics

To evaluate the accuracy of side-chain modeling methods, we employ the percentage of correct predictions using a tolerance criterion of 20° for all side-chain dihedral angles ranging from χ_1_ to χ_4_ (denote as Accuracy). The Mean Absolute Error (MAE (χ)) quantifies the average absolute difference between the native and predicted values for each side-chain dihedral angle. The Root Mean Square Deviation (RMSD) is computed over non-hydrogen side-chain atoms, excluding Ala and Gly residues. When determining RMSD and MAE (χ), we account for the symmetry of residues such as Asp, Glu, Phe, Arg, and Tyr ^3, 10^. A residue is defined as a core residue if there are more than 20 residues where the C_β_-C_β_ distance (C_α_ for Gly) is within 10 Å.

For assessing molecular docking performance, we initially align AlphaFold2’s predicted backbones with the experimentally determined ones. We then use two criteria for evaluation. The stricter criterion defines successful docking cases as those that reproduce experimental coordinates of the ligands within an RMSD of 2 Å. This is assessed for the top 1, top 3, and top 5 poses. Since the predicted and experimental backbones are not strictly aligned, we also adopt a more lenient criterion, which defines successful docking cases as those reproducing experimental coordinates of the ligands within an RMSD of 3 Å. Additionally, we compute the average RMSD for the top 1 pose to provide a comparative metric.

### Data and Software Availability

The code and pre-trained models of OPUS-Rota5 as well as the datasets used in the study can be downloaded from http://github.com/OPUS-MaLab/opus_rota5. They are freely available for academic usage only.

## Results

### Side-chain modeling performance of different methods

We evaluate the performance of OPUS-Rota5 in side-chain modeling against two sampling-based methods (OSCAR-star and RosettaPacker) and three deep learning-based methods (DLPacker, AttnPacker, and our preceding method, OPUS-Rota4) across three native backbone test sets (CAMEO65, CASP15, and CAMEO82). OPUS-Rota5 consistently surpasses the other methods in various metrics, including the percentage of correct prediction, the mean absolute error of each dihedral angle, and side-chain RMSD, whether measured across all residues (Table 1) or solely within core residues (SI appendix, Table S1). Additionally, the percentage of correct prediction for each residue type on each test set is detailed in SI appendix, Table S2 (all residues) and SI appendix, Table S3 (core residues).

**Table 1.**
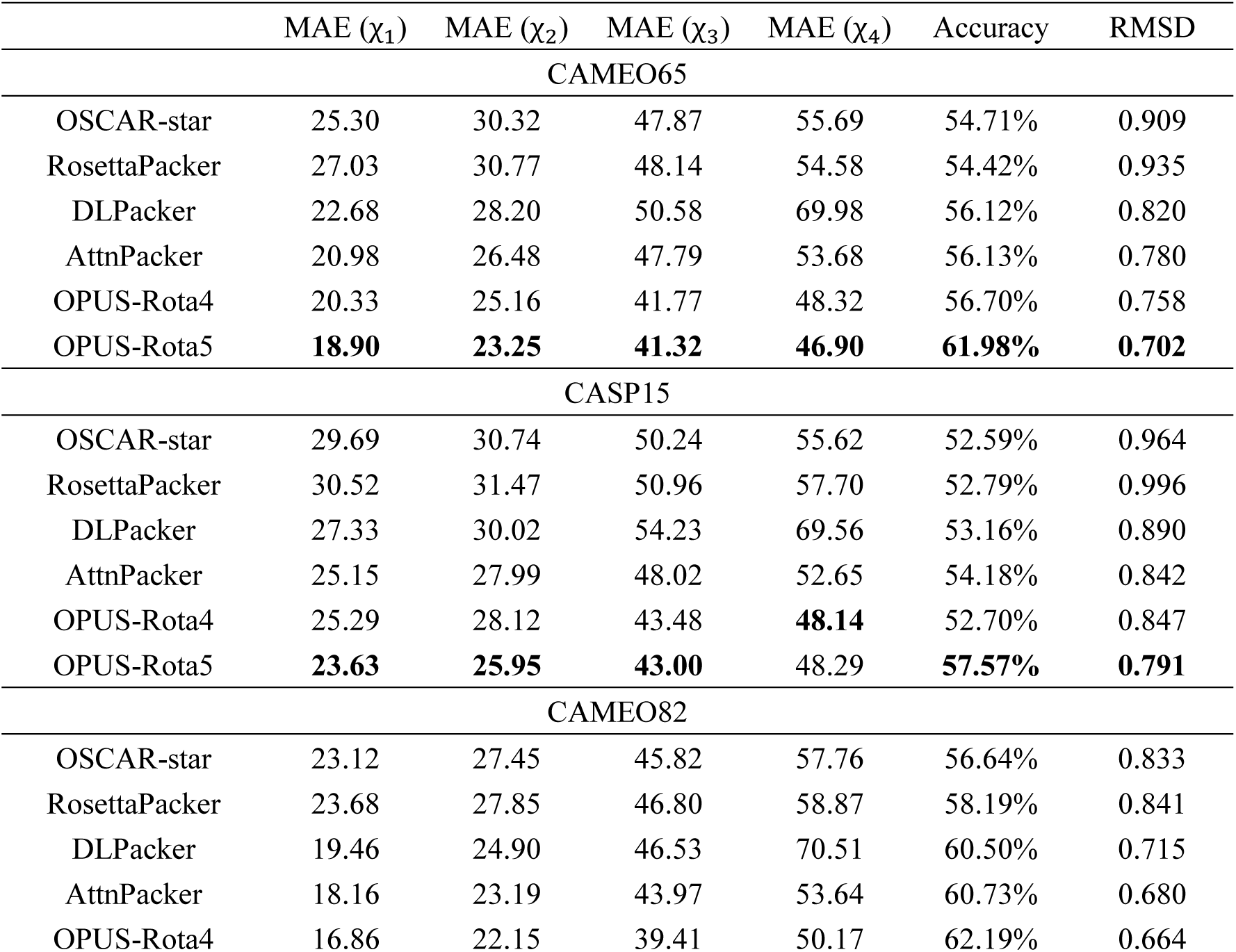
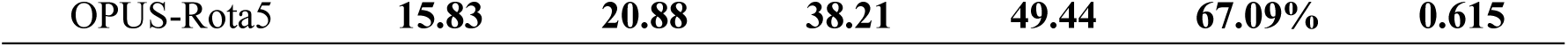
The side-chain modeling results of various methods measured by all residues. Best results for each metric are shown in boldface.

For the side-chain modeling method DiffPacker, it relies on the given C_β_ atom as inputs, which is different from the aforementioned methods. To ensure a fair comparison, we reconstruct the C_β_ atom using RosettaPacker and evaluate the performance of DiffPacker. As indicated in SI appendix, Table S4, DiffPacker achieves side-chain modeling results comparable to OPUS-Rota5 when employing the native C_β_ atom as inputs. However, its performance notably diminishes when using the predicted C_β_ atom as inputs. Across the targets of all three native backbone test sets, the percentage of correct prediction, measured by core residues, dips from 85.73% to 79.32%, while that for OPUS-Rota5 is 86.13%. This observation underscores the criticality of accurately positioning the C_β_ atom in side-chain modeling and it should be an important issue for future study.

In Figure 2, we present several examples of side-chain modeling results from different methods. The results reveal that the side chains predicted by OPUS-Rota5 are significantly closer to their native counterparts comparing to those predicted by other state-of-the-art methods.

**Figure 2.**
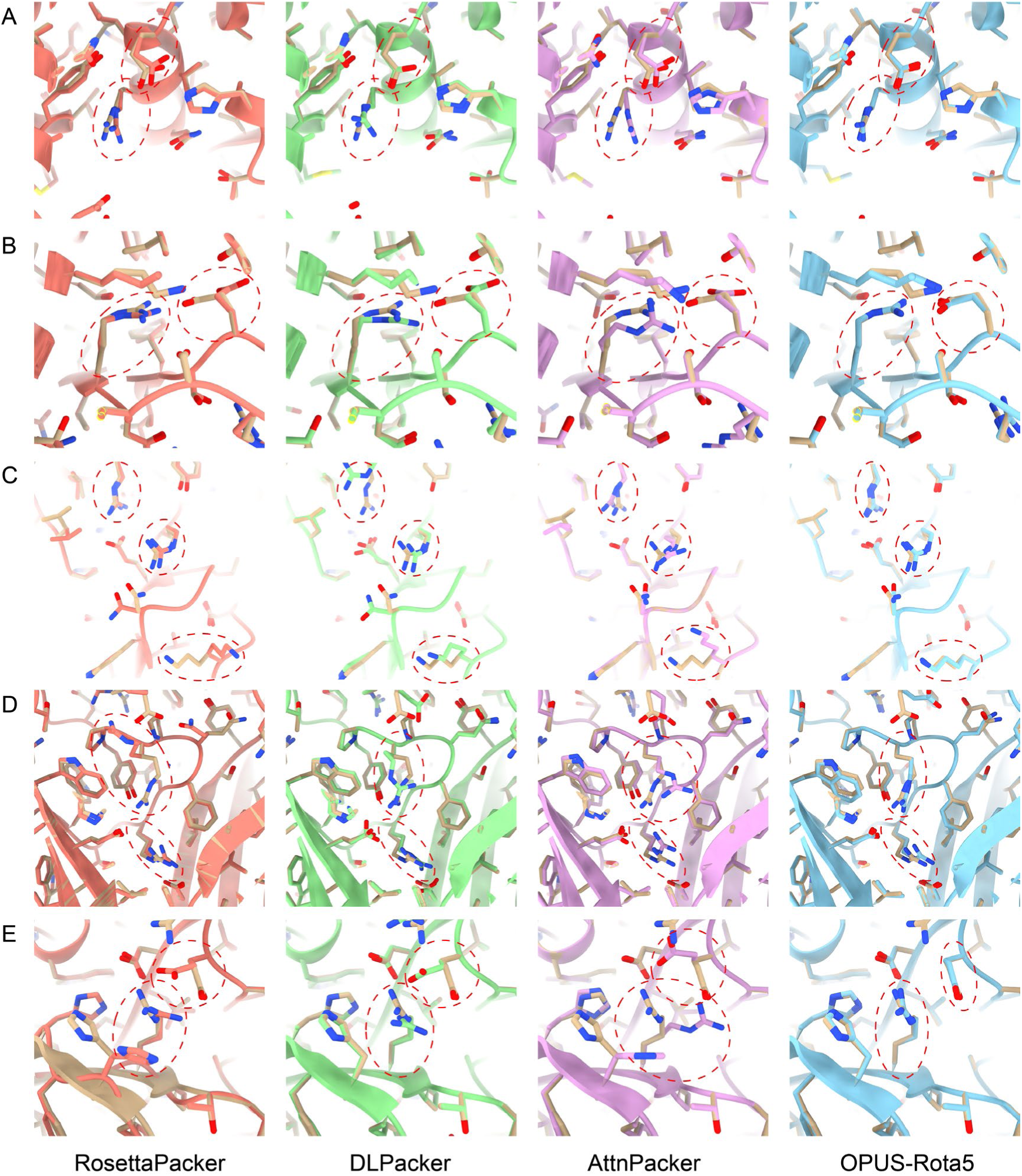
Side-chain predictions compared with experimental structures. Experimental structures are colored in gold. Predictions from RosettaPacker, DLPacker, AttnPacker, and OPUS-Rota5 are colored in orange, green, violet, and blue, respectively. A) Comparison around residues 322∼329 on T1180-D1. B) Comparison around residues 226∼229 on T1129s2-D1. C) Comparison around residues 58∼64 of Chain A on 2023-06-24_000000193_1. D) Comparison around residues 27∼35 of Chain A on 2023-07-01_00000022_1. E) Comparison around residues 376∼378 of Chain B on 2021-10-02-00000013_1.

### Ablation study of OPUS-Rota5

Compared to OPUS-Rota4, OPUS-Rota5 omits the time-consuming optimization procedure and modifies the architectures of the feature extraction and aggregation modules. Specifically, we employ a modified 3D-Unet to capture the local environmental features of each residue and subsequently utilize RotaFormer module to aggregate various types of features. Additionally, we incorporate four BiLSTM layers to further aggregate sequence features from RotaFormer. Self-attention is pivotal for capturing long-term information since two residues might be proximate in 3D space even though they may be relatively distant in 1D sequence. Conversely, LSTM effectively captures short-term information, which is also vital since side-chain conformations are largely determined by their immediate surroundings. As illustrated in Figure S1, all of these modifications contribute to the final accuracy.

Moreover, we implement a 3D-Transformer module for local environmental feature extraction by substituting the residue blocks with EPA blocks from UNETR++ ^50^. In addition, we introduce the protein evolutionary features from ESM-1b ^51^ as one of the sources of the sequence features for training the RotaFormer module. However, both of these modifications cannot improve the side-chain modeling performance of OPUS-Rota5 (Figure S1).

### Applications of OPUS-Rota5 in molecular docking

To evaluate the efficacy of utilizing side chains reconstructed by OPUS-Rota5 in molecular docking, we compare the docking results using the original predicted structures from AlphaFold2 (AF2 in Table 2) and the structures with the backbones from AlphaFold2 while side chains reconstructed by OPUS-Rota5 (AF2+Rota5 in Table 2). We employ a set of 25 complexes, involving G protein-coupled receptors (GPCRs) and their native small molecule ligands, as the docking targets. Two docking programs, AutoDock-Vina and Dock6.10, are utilized in this study. For AutoDock- Vina, we set the box size as (30, 30, 30) and set the exhaustiveness to 16. For Dock6.10, we proceed with the default parameters for docking.

**Table 2.**
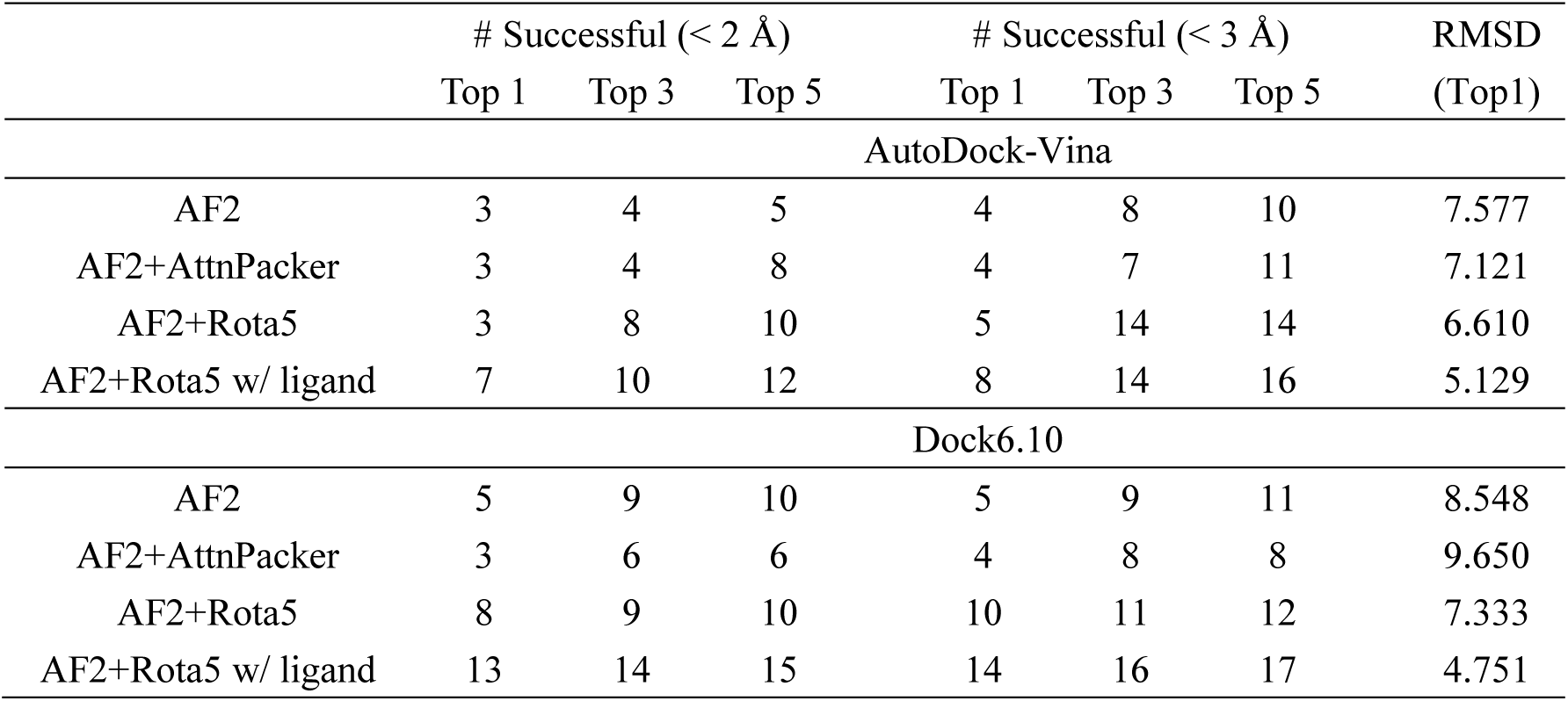
The docking results conducted by AutoDock-Vina and Dock6.10 on 25 GPCR targets with different side-chain conformations.

|  | # Successful (< 2 Å) |  |  | # Successful (< 3 Å) |  |  | RMSD |
| --- | --- | --- | --- | --- | --- | --- | --- |
|  | Top 1 | Top 3 | Top 5 | Top 1 | Top 3 | Top 5 | (Top1) |
| AutoDock-Vina |  |  |  |  |  |  |  |
| AF2 | 3 | 4 | 5 | 4 | 8 | 10 | 7.577 |
| AF2+AttnPacker | 3 | 4 | 8 | 4 | 7 | 11 | 7.121 |
| AF2+Rota5 | 3 | 8 | 10 | 5 | 14 | 14 | 6.610 |
| AF2+Rota5 w/ ligand | 7 | 10 | 12 | 8 | 14 | 16 | 5.129 |
| Dock6.10 |  |  |  |  |  |  |  |
| AF2 | 5 | 9 | 10 | 5 | 9 | 11 | 8.548 |
| AF2+AttnPacker | 3 | 6 | 6 | 4 | 8 | 8 | 9.650 |
| AF2+Rota5 | 8 | 9 | 10 | 10 | 11 | 12 | 7.333 |
| AF2+Rota5 w/ ligand | 13 | 14 | 15 | 14 | 16 | 17 | 4.751 |

As indicated in Table 2, the success rates of reproducing experimental coordinates of the ligands, both within a stricter 2 Å RMSD criterion and a more lenient 3 Å RMSD criterion, as well as the average RMSD for the top 1 pose, exhibit improvement after side-chain reconstruction with OPUS-Rota5 when utilizing both AutoDock-Vina and Dock6.10, respectively. Consequently, OPUS-Rota5 emerges as a valuable tool for molecular docking, especially for targets that lack experimentally determined structures but possess relatively accurate predicted backbones (e.g. high-homology targets).

In Figure 3, we present several examples of Dock6.10’s docking results on the structures predicted by AlphaFold2 (AF2) and the structures with the same backbones while side chains reconstructed by OPUS-Rota5 (AF2+Rota5). The results underscore AlphaFold2’s ability to provide relatively accurate backbone structures around the ligand binding site, an important foundation for side-chain modeling. After side-chain reconstruction, AF2+Rota5 demonstrates enhanced docking performance, identifying the ligand poses closer to the native results. Moreover, as depicted in Figure 3B, the RMSD of the top 1 pose for AF2+Rota5 is 0.914, closely paralleling the third pose of AF2, which is 0.902. However, their grid scores, calculated by Dock6.10, diverge. The grid score, an approximation of the molecular mechanics’ energy function considering only through space interactions, remarks the stability of protein-ligand complex, the lower the score, the more stable the complex. The grid score is -52.60 for AF2+Rota5 and -30.83 for AF2, which indicates that the ligand-protein binding pose based on AF2+Rota5 is more favorable. This observation is further corroborated by Figure 3C and Figure 3D. For instance, in Figure 3D, the grid score for the top 1 pose is -33.81 for AF2+Rota5 and -30.25 for AF2.

**Figure 3.**
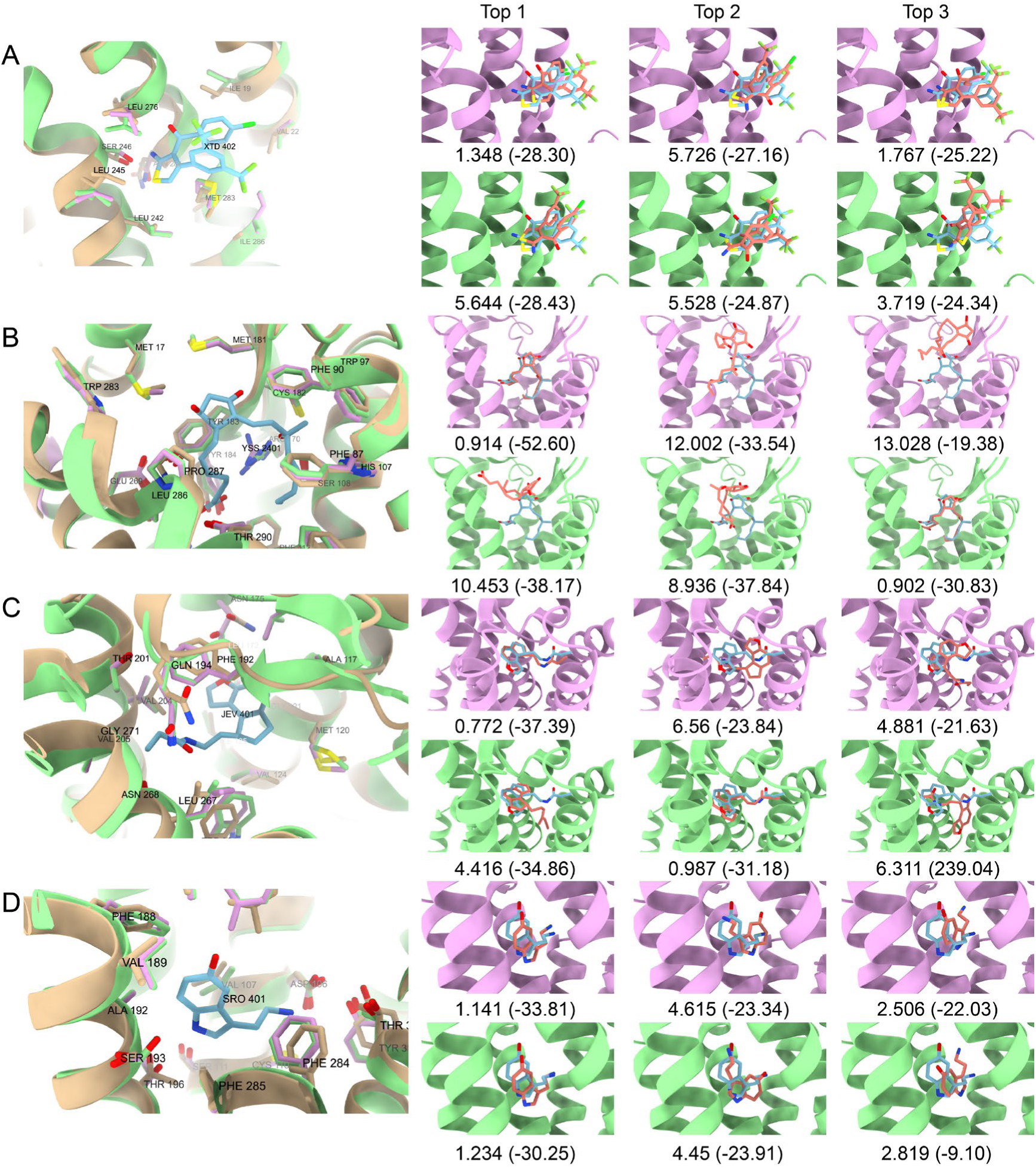
Docking results from Dock6.10 on original AlphaFold2 predicted structures (AF2) and structures with same backbone while side chains reconstructed by OPUS-Rota5 (AF2+Rota5). Experimental protein and ligand structures are colored in gold and blue, respectively. Protein structures predicted by AlphaFold2 are in green, while side chains reconstructed by OPUS-Rota5 are in pink. On each panel’s left side, the local environment of the ligand binding site is displayed. On the right, the top 3 docking poses on AF2+Rota5 (pink structures, top) and AF2 (green structures, bottom) are shown. Numbers outside parentheses represent RMSDs, while numbers inside are grid scores from Dock6.10. Panels A-D display docking results on 7DL3, 7M8W, 7VH0, and 7YS6, respectively.

When we represent the ligand as corresponding atoms in 3D grids, OPUS-Rota5 can capture the ligand information through its modified 3D-Unet module. Therefore, we place the ligand at the experimentally determined position and reconstruct the side chains for the AlphaFold2 predicted backbone. The results (AF2+Rota5 w/ ligand in Table 2) demonstrate that, with the help of ligand information, the reconstructed side chains are more conducive to yielding the correct docking pose through the docking program. This suggests that OPUS-Rota5 is sensitive to changes in the local environment and may prove useful for flexible docking and ligand design.

Figure 4 illustrates several examples of AutoDock-Vina’s docking results on AlphaFold2’s predicted backbones with side chains reconstructed by OPUS-Rota5, both with ligand information (AF2+Rota5 w/ ligand) and without ligand information (AF2+Rota5). Assisted by ligand information, OPUS-Rota5 can deliver side chains that are closer to the native states, such as GLN134 in Figure 4A and PHE210 in Figure 4B. As depicted in Figure 4C, for the top 1 pose, the affinity calculated by AutoDock-Vina for AF2+Rota5 w/ ligand is -9.5, compared to -8.7 for AF2+Rota5. This indicates that the ligand-protein binding pose based on AF2+Rota5 w/ ligand is more favorable, even though their RMSD values are close (1.640 for AF2+Rota5 w/ ligand and 1.722 for AF2+Rota5).

**Figure 4.**
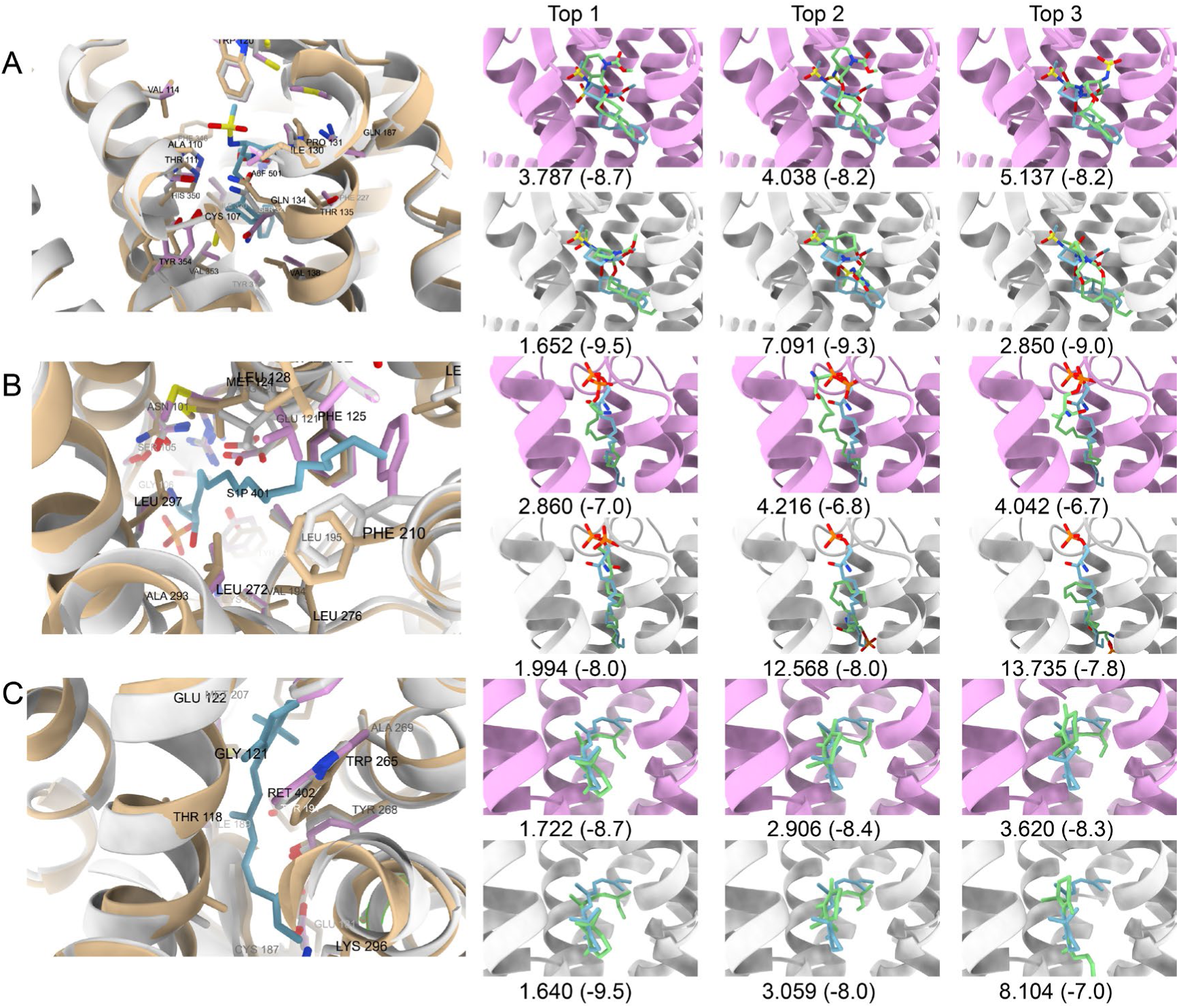
AutoDock-Vina docking results on AlphaFold2’s predicted backbones with side chains reconstructed by OPUS-Rota5, with and without ligand information (AF2+Rota5 w/ ligand and AF2+Rota5, respectively). Experimental protein and ligand structures are colored in gold and blue, respectively. Side chains reconstructed by OPUS-Rota5 are colored in pink without ligand information, and in gray with ligand information. On the left side of each panel, the local environment of the ligand binding site is displayed. On the right, the top 3 docking poses on AF2+Rota5 (pink structures, top) and AF2+Rota5 w/ ligand (gray structures, bottom) are shown. Numbers outside parentheses represent RMSDs, while numbers inside are affinities from AutoDock-Vina. Panels A-C display docking results on 7SR8, 7VIE, and 7ZBC, respectively

## Concluding Discussion

In this study, we introduce OPUS-Rota5, a two-stage protein side-chain modeling method that utilizes a modified 3D-Unet to capture the local environmental features of each residue. Subsequently, it employs RotaFormer module to aggregate various types of features. Our evaluations on three native test sets underscore the superiority of OPUS-Rota5 over several other state-of-the-art side-chain modeling methods. For instance, when tested on the recently released targets (CAMEO82), OPUS-Rota5 achieves a percentage of correct predictions of 67.09% using a tolerance criterion of 20° for all side-chain dihedral angles (from χ_1_to χ_4_), measured by all residues. In comparison, OPUS-Rota4 and AttnPacker achieve 62.19% and 60.73%, respectively. These results suggest that OPUS-Rota5 possesses commendable generalizability, establishing it as a valuable side-chain modeling tool for the scientific community.

While the positions of C_β_ atoms are typically determined through backbone atoms, findings from this study highlight the critical role of the accurate positioning of the C_β_ atom in side-chain modeling. Consequently, future research directions may necessitate a focus on the precise prediction of C_β_ atom positions.

For a target without an experimentally determined structure, if molecular docking is intended, AlphaFold2 can be utilized first to provide a predicted structure. However, from recent insights into GPCR targets, its predicted side chains of certain residues are not accurate enough for the purpose of successful docking. In such cases, OPUS-Rota5 can be employed to reconstruct the side chains based on the backbones from AlphaFold2. Our findings, as shown in docking results by both AutoDock-Vina and Dock6.10, illustrate the side chains reconstructed by OPUS-Rota5 enable higher success rates of docking. This demonstrates that OPUS-Rota5 adeptly rearranges the side chains for AlphaFold2’s predicted structures, thereby optimizes molecular docking. Consequently, it emerges as a valuable tool for molecular docking, especially for targets lacking experimentally determined structures but possessing relatively accurate predicted backbones, such as high-homology targets.

Moreover, OPUS-Rota5 can incorporate ligand information through its modified 3D-Unet module, enabling it to render side chains influenced by the ligand. The results showcase that, with the aid of ligand information, the reconstructed side chains facilitate a more accurate docking pose through the docking program compared to those without ligand information. Additionally, the results indicate that, when assisted by ligand information, OPUS-Rota5 can produce side chains that more closely resemble the native states of protein-ligand complexes. The sensitivity of OPUS- Rota5 to alterations in the local environment may be advantageous for flexible docking and ligand design.

## Acknowledgements

Jianpeng Ma wants to thank the supports from National Key Research and Development Program of China (No. 2021YFF1200400), Shanghai Municipal Science and Technology Major Project (No.2018SHZDZX01), and ZJLab. Gang Xu wants to thank the support from National Natural Science Foundation of China (No. 32300535).

## Notes

### Competing Interest Statement

The authors have declared no competing interest.

