## Supplementary material for "OPUS-Rota5: A Highly Accurate Protein Side-chain Modeling Method with 3D-Unet and RotaFormer": SI

Xu et al.

**Table S1**. The side-chain modeling results of various methods measured by core residues. Best results for each metric are shown in boldface.

| | MAE ($\chi_1$) | MAE ($\chi_2$) | MAE ($\chi_3$) | MAE ($\chi_4$) | Accuracy | RMSD |
|---|---|---|---|---|---|---|
| | | | CAMEO65 | | | |
| OSCAR-star | 14.90 | 18.61 | 37.83 | 34.85 | 73.36% | 0.535 |
| RosettaPacker | 14.21 | 17.63 | 33.83 | 39.07 | 76.23% | 0.484 |
| DLPacker | 16.48 | 15.85 | 33.85 | 35.30 | 75.29% | 0.489 |
| AttnPacker | 11.49 | 16.09 | 36.47 | 39.64 | 77.06% | 0.410 |
| OPUS-Rota4 | 11.32 | 14.11 | 27.39 | 27.51 | 78.18% | 0.396 |
| OPUS-Rota5 | **9.50** | **11.93** | **22.34** | **24.52** | **83.77%** | **0.338** |
| | | | CASP15 | | | |
| OSCAR-star | 16.87 | 20.19 | 25.98 | 33.06 | 72.10% | 0.542 |
| RosettaPacker | 16.33 | 20.95 | 26.36 | 39.98 | 75.61% | 0.525 |
| DLPacker | 14.21 | 17.75 | 33.49 | 36.94 | 77.03% | 0.468 |
| AttnPacker | 13.53 | 16.30 | 27.90 | 35.12 | 75.87% | 0.439 |
| OPUS-Rota4 | 10.92 | 15.72 | **20.31** | 29.55 | 77.38% | 0.393 |
| OPUS-Rota5 | **9.67** | **12.78** | 20.68 | **27.43** | **83.43%** | **0.341** |
| | | | CAMEO82 | | | |
| OSCAR-star | 14.68 | 17.89 | 39.15 | 34.41 | 73.91% | 0.517 |
| RosettaPacker | 14.06 | 18.33 | 35.38 | 35.99 | 76.84% | 0.494 |
| DLPacker | 10.38 | 14.61 | 34.72 | 37.87 | 80.54% | 0.377 |
| AttnPacker | 9.90 | 13.31 | 37.31 | 32.74 | 80.06% | 0.372 |
| OPUS-Rota4 | 7.73 | 12.11 | 29.21 | 24.72 | 83.44% | 0.330 |
| OPUS-Rota5 | **7.40** | **10.31** | **26.87** | **23.45** | **87.54%** | **0.292** |

**Table S2**. The percentage of correct prediction for each residue type measured by all residues. Best results for each metric are shown in boldface.

| | OSCAR-star | RosettaPacker | DLPacker | AttnPacker | OPUS-Rota4 | OPUS-Rota5 |
|---|---|---|---|---|---|---|
| CAMEO65 | | | | | | |
| C | 81.06% | 76.14% | 83.71% | 84.09% | 83.71% | **89.02%** |
| D | 49.35% | 50.55% | 56.92% | 56.46% | 57.84% | **62.08%** |
| E | 23.68% | 22.46% | 19.86% | 25.52% | 24.98% | **33.08%** |
| F | 68.97% | 74.43% | 79.63% | 80.69% | 75.10% | **85.89%** |
| H | 50.67% | 57.59% | 66.29% | 62.50% | 65.63% | **72.54%** |
| I | 68.66% | 69.31% | 67.29% | 68.50% | 71.16% | **74.96%** |
| K | 18.80% | 17.60% | 9.18% | 14.42% | 18.03% | **22.23%** |
| L | 72.53% | 71.57% | 75.41% | 74.73% | 75.80% | **79.24%** |
| M | 36.16% | 36.25% | 32.92% | 25.94% | 40.90% | **46.13%** |
| N | 45.45% | 48.45% | 53.44% | 50.00% | 53.22% | **61.09%** |
| P | 76.02% | 77.58% | 77.46% | 76.74% | 73.98% | **79.98%** |
| Q | 25.40% | 28.47% | 25.40% | 26.57% | 32.12% | **38.69%** |
| R | 21.72% | 19.81% | 16.95% | 10.81% | 25.53% | **32.84%** |
| S | 61.91% | 59.39% | 66.96% | 67.65% | 63.48% | **71.65%** |
| T | 79.58% | 79.58% | 80.15% | 78.24% | 79.87% | **83.70%** |
| V | 87.77% | 88.62% | 89.71% | 88.70% | 88.95% | **90.05%** |
| W | 64.55% | 73.58% | 83.95% | 78.60% | 88.96% | **91.30%** |
| Y | 71.84% | 77.63% | 85.39% | 83.56% | 78.39% | **89.35%** |
| CASP15 | | | | | | |
| C | 74.23% | 68.04% | 72.16% | 76.29% | 71.13% | **83.51%** |
| D | 43.48% | 43.30% | 45.29% | **49.28%** | 46.38% | 49.09% |
| E | 22.59% | 22.26% | 15.28% | 22.26% | 23.92% | **30.73%** |
| F | 62.37% | 68.56% | 72.68% | 75.00% | 71.91% | **76.03%** |
| H | 50.60% | 51.20% | 55.42% | 49.40% | 53.61% | **59.64%** |
| I | 71.51% | 67.51% | 68.78% | 69.69% | 71.32% | **76.04%** |
| K | 12.34% | 11.69% | 8.66% | 10.39% | 13.20% | **16.23%** |
| L | 64.36% | 63.87% | 67.87% | **69.73%** | 64.94% | 67.77% |
| M | 32.43% | 34.05% | 25.95% | 23.24% | 36.22% | **40.00%** |
| N | 42.33% | 45.36% | 48.81% | 48.81% | 45.36% | **52.48%** |
| P | 73.33% | 72.95% | 71.43% | 69.52% | 67.05% | **71.81%** |
| Q | 23.31% | 23.60% | 19.94% | 26.69% | 31.18% | **32.30%** |
| R | 13.32% | 12.91% | 13.93% | 6.97% | 18.24% | **21.93%** |
| S | 51.65% | 50.83% | 52.64% | 53.30% | 46.37% | **59.24%** |
| T | 77.55% | 78.33% | 77.55% | **78.02%** | 76.63% | 77.86% |
| V | 89.69% | 89.55% | 89.27% | 91.10% | 88.42% | **92.80%** |
| W | 60.58% | 71.53% | 79.56% | 69.34% | 82.48% | **84.67%** |
| Y | 68.75% | 73.81% | 81.85% | 80.95% | 71.73% | **81.85%** |
| CAMEO82 | | | | | | |

| | | | | | | |
|---|---|---|---|---|---|---|
| C | 81.82% | 74.70% | 85.77% | 86.56% | 84.98% | **92.89%** |
| D | 50.60% | 52.01% | 59.22% | 61.13% | 61.91% | **67.14%** |
| E | 23.03% | 24.23% | 21.69% | 26.51% | 28.58% | **35.68%** |
| F | 71.04% | 80.52% | 88.48% | 87.08% | 81.92% | **91.28%** |
| H | 53.70% | 58.26% | 69.13% | 62.17% | 66.09% | **78.48%** |
| I | 75.19% | 74.66% | 75.19% | 75.71% | 79.99% | **82.23%** |
| K | 15.93% | 14.57% | 8.79% | 12.93% | 13.79% | **17.79%** |
| L | 78.27% | 76.99% | 81.58% | 82.62% | 81.58% | **84.49%** |
| M | 37.16% | 37.79% | 34.86% | 27.14% | 45.51% | **50.10%** |
| N | 51.19% | 55.15% | 60.69% | 57.23% | 62.48% | **68.81%** |
| P | 76.98% | 77.75% | 79.27% | 77.55% | 78.03% | **81.66%** |
| Q | 22.09% | 30.93% | 28.14% | 27.44% | 36.74% | **42.44%** |
| R | 21.61% | 20.04% | 18.09% | 11.87% | 28.76% | **32.56%** |
| S | 63.72% | 63.50% | 69.68% | 68.06% | 67.99% | **74.69%** |
| T | 83.12% | 83.20% | 83.75% | 85.57% | 85.02% | **88.17%** |
| V | 91.11% | 92.48% | 94.12% | 93.20% | 93.14% | **95.03%** |
| W | 68.97% | 75.86% | 85.06% | 87.64% | 90.80% | **91.38%** |
| Y | 67.47% | 78.83% | 83.16% | 85.46% | 82.02% | **90.94%** |

**Table S3**. The percentage of correct prediction for each residue type measured by core residues. Best results for each metric are shown in boldface.

| | OSCAR-star | RosettaPacker | DLPacker | AttnPacker | OPUS-Rota4 | OPUS-Rota5 |
|---|---|---|---|---|---|---|
| CAMEO65 | | | | | | |
| C | 79.31% | 78.62% | 84.83% | 85.52% | 87.59% | **88.97%** |
| D | 56.55% | 66.90% | 73.10% | 67.59% | 77.24% | **82.07%** |
| E | 33.60% | 39.20% | 29.60% | 28.00% | 40.00% | **52.00%** |
| F | 73.01% | 82.39% | 86.93% | 88.07% | 84.38% | **93.47%** |
| H | 57.50% | 70.83% | 78.33% | 70.00% | 81.67% | **86.67%** |
| I | 75.86% | 78.98% | 75.70% | 77.67% | 81.28% | **84.89%** |
| K | 43.27% | 41.35% | 22.12% | 25.96% | 42.31% | **52.88%** |
| L | 78.62% | 78.39% | 84.81% | 82.59% | 86.57% | **89.25%** |
| M | 42.78% | 48.66% | 41.71% | 35.83% | 54.01% | **59.89%** |
| N | 55.37% | 66.10% | 63.28% | 56.50% | 71.19% | **79.10%** |
| P | 82.58% | 81.46% | 84.27% | 79.21% | 81.46% | **87.08%** |
| Q | 33.09% | 41.18% | 36.76% | 38.24% | 52.94% | **63.24%** |
| R | 30.94% | 34.53% | 30.22% | 16.55% | 43.88% | **58.99%** |
| S | 70.50% | 69.88% | 76.40% | 75.47% | 74.84% | **80.75%** |
| T | 86.94% | 87.31% | 87.69% | 86.57% | 89.18% | **93.28%** |
| V | 90.48% | 92.73% | 91.87% | 90.83% | 92.39% | **92.56%** |
| W | 69.70% | 84.09% | 87.12% | 83.33% | 93.18% | **93.94%** |
| Y | 74.44% | 85.93% | 92.96% | 88.15% | 86.30% | **95.93%** |
| CASP15 | | | | | | |
| C | 85.29% | 82.35% | 85.29% | 88.24% | 85.29% | **94.12%** |
| D | 57.14% | 67.35% | 77.55% | 71.43% | **83.67%** | 79.59% |
| E | 31.82% | 38.64% | 25.00% | 43.18% | 50.00% | **59.09%** |
| F | 71.88% | 86.72% | 85.94% | 87.50% | 89.06% | **92.97%** |
| H | 62.16% | 70.27% | 81.08% | 70.27% | **83.78%** | 81.08% |
| I | 76.67% | 72.86% | 77.62% | 77.62% | 82.86% | **86.67%** |
| K | 21.57% | 31.37% | 17.65% | 17.65% | 25.49% | **35.29%** |
| L | 74.70% | 71.43% | 78.27% | 80.36% | 82.14% | **83.63%** |
| M | 43.66% | 49.30% | 35.21% | 30.99% | 52.11% | **54.93%** |
| N | 52.27% | 59.09% | 63.64% | 56.82% | 63.64% | **72.73%** |
| P | 70.31% | 76.56% | 73.44% | 67.19% | 71.88% | **75.00%** |
| Q | 47.06% | 52.94% | 35.29% | 38.24% | **58.82%** | 55.88% |
| R | 22.22% | 25.93% | 25.93% | 9.26% | 44.44% | **46.30%** |
| S | 61.21% | 64.66% | 68.10% | 69.83% | 64.66% | **83.62%** |
| T | 84.62% | 86.32% | 88.89% | 87.18% | **88.03%** | 87.18% |
| V | 92.36% | 91.27% | 93.45% | 93.45% | 94.18% | **97.09%** |
| W | 68.63% | 84.31% | 88.24% | 80.39% | 92.16% | **96.08%** |
| Y | 73.44% | 85.94% | 94.53% | 89.06% | 82.81% | **95.31%** |
| CAMEO82 | | | | | | |

| | | | | | | |
|---|---|---|---|---|---|---|
| C | 88.15% | 84.44% | 94.07% | 91.85% | 94.81% | **97.04%** |
| D | 62.38% | 62.87% | 71.29% | 74.26% | 80.20% | **83.66%** |
| E | 38.31% | 52.60% | 32.47% | 34.42% | 43.51% | **54.55%** |
| F | 72.39% | 85.69% | 91.82% | 89.78% | 87.32% | **94.89%** |
| H | 56.16% | 63.70% | 81.51% | 66.44% | 77.40% | **90.41%** |
| I | 81.07% | 79.62% | 81.65% | 81.21% | 86.85% | **87.86%** |
| K | 30.36% | 27.68% | 18.75% | 22.32% | 33.93% | **45.54%** |
| L | 82.51% | 81.21% | 86.83% | 86.53% | 87.24% | **89.55%** |
| M | 42.92% | 45.83% | 42.92% | 35.42% | 55.42% | **60.00%** |
| N | 60.50% | 70.00% | 71.50% | 66.00% | 82.00% | **84.00%** |
| P | 81.35% | 84.52% | 85.32% | 82.54% | 84.52% | **84.92%** |
| Q | 30.94% | 52.52% | 38.85% | 36.69% | 61.15% | **66.91%** |
| R | 37.00% | 35.50% | 31.50% | 17.50% | 51.50% | **60.00%** |
| S | 70.27% | 76.22% | 77.84% | 74.05% | 80.27% | **87.30%** |
| T | 87.35% | 89.76% | 90.96% | 90.36% | 93.07% | **94.58%** |
| V | 94.21% | 95.32% | 96.43% | 95.81% | 96.67% | **98.03%** |
| W | 73.65% | 86.49% | 92.57% | 89.86% | 96.62% | **98.65%** |
| Y | 70.26% | 84.55% | 90.96% | 89.21% | 89.21% | **95.63%** |

**Table S4**. The side-chain modeling performance of DiffPacker using experimental $C_\beta$ atoms (DiffPacker (real $C_\beta$)) and predicted $C_\beta$ atoms by RosettaPacker (DiffPakcer (pred $C_\beta$)) on targets across all three native backbone test sets. Best results for each metric are shown in boldface.

| | MAE ($\chi_1$) | MAE ($\chi_2$) | MAE ($\chi_3$) | MAE ($\chi_4$) | Accuracy | RMSD |
|---|---|---|---|---|---|---|
| All residues | | | | | | |
| DiffPakcer (pred $C_\beta$) | 24.80 | 28.45 | 47.48 | 59.50 | 58.71% | 0.865 |
| DiffPacker (real $C_\beta$) | **17.34** | **22.15** | 40.81 | 54.96 | **65.79%** | **0.655** |
| OPUS-Rota5 | 18.51 | 22.57 | **40.40** | **48.47** | 63.51% | 0.680 |
| Core residues | | | | | | |
| DiffPakcer (pred $C_\beta$) | 13.73 | 16.21 | 35.34 | 34.31 | 79.32% | 0.454 |
| DiffPacker (real $C_\beta$) | 9.44 | 11.54 | 25.96 | 29.92 | 85.73% | 0.311 |
| OPUS-Rota5 | **8.45** | **10.87** | **23.16** | **24.53** | **86.13%** | **0.309** |

**Figure S1**. The percentage of correct predictions, using a tolerance criterion of 20° for all side-chain dihedral angles, for different versions of OPUS-Rota5, measured by all residues on CAMEO65.

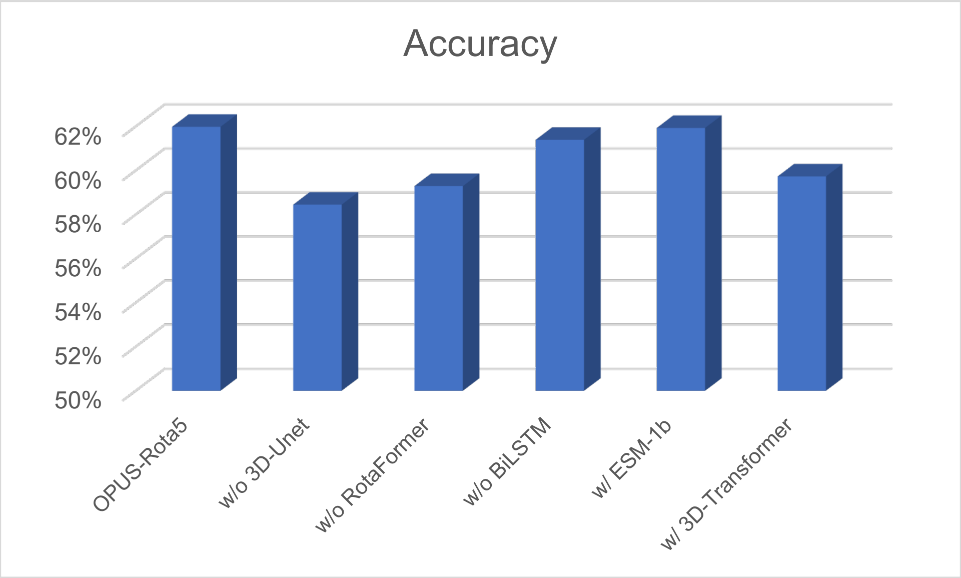
